# Distinct DNA Methylation Signatures in Maternal Blood Reveal Unique Immune Cell Shifts in Preeclampsia and the Pregnancy-Postpartum Transition

**DOI:** 10.1101/2024.12.13.628167

**Authors:** Laiba Jamshed, Keaton W Smith, Samantha L Wilson

## Abstract

Preeclampsia (PE) is a hypertensive disorder of pregnancy characterized by immune dysregulation and significant risks to maternal and fetal health. While current management relies on high-risk patient monitoring and early diagnosis, these methods are costly and burdensome, especially for low-risk pregnancies. There is a pressing need for non-invasive tools to predict and monitor PE. DNA methylation (DNAm) is a type of DNA modification that influences gene expression, and has been associated with immune cell dynamics and PE pathogenesis. This study explores whether DNAm-based immune cell composition profiling can provide insights into PE-related immune dysregulation. We conducted a search in the Gene Expression Omnibus (GEO) for DNA methylation datasets using Illumina 27K, 450K, and EPIC arrays from maternal blood in both healthy and PE pregnancies. We found two studies that met our criteria, involving a total of 24 healthy pregnancies and 14 with PE. To estimate the composition of immune cells (including CD8T, CD4T, Monocytes, Natural Killer, Neutrophils, Eosinophils, and B cells) based on DNA methylation data, we employed the R package EpiDISH. We used a linear model to compare statistical differences in the proportions of immune cells between PE cases and the control group. Longitudinal trends were also examined to capture immune cell shifts from pregnancy to postpartum. We found that monocyte proportions were significantly reduced in preeclamptic pregnancies compared to normotensive pregnancies (p=0.013). No significant differences were observed in other immune cell types, including T cells, B cells, neutrophils, eosinophils, and natural killer cells. Longitudinal analyses revealed substantial immune cell shifts in the postpartum period, including increased monocytes, B cells, CD4+ T cells, and CD8+ T cells, emphasizing the importance of gestational age in immune dynamics. These findings support DNAm profiling as a valuable tool for understanding immune cell dynamics in PE. Reduced monocyte proportions in PE highlight the role of immune dysregulation in its pathogenesis. Longitudinal sampling provides additional insights into the evolution of immune changes throughout pregnancy and postpartum, offering potential for developing predictive and monitoring tools for PE. Future studies with larger, more diverse cohorts are essential to refine the utility of DNAm in pregnancy complications.

## 2.0 Introduction

### 2.1 Overview of Preeclampsia

Preeclampsia (PE) is a pregnancy-specific disorder characterized by new-onset hypertension and proteinuria after 20 weeks of gestation.^1^ Occurring in an estimated 3–5% of pregnancies, PE is a leading cause of both maternal and fetal morbidity and mortality worldwide.^2^ Although PE is a common complication of pregnancy, its exact etiology and pathogenesis are not fully understood, making early detection and intervention challenging. As a multisystemic disorder, the pathogenesis of PE has been associated with poor placentation, placental hypoxia, endothelial dysfunction, impaired angiogenesis, and excessive maternal inflammation.^2,3^ Together, these altered pathways ultimately lead to a placental ischemic microenvironment, inadequate uterine vascular remodeling, compromised blood perfusion and oxidative stress.^2–5^ Collectively, these factors contribute to the hallmark clinical features of hypertension and organ dysfunction including renal, hepatic, hematological or neurological complications.^2–5^ If left untreated, PE can lead to severe maternal complications, including cerebral hemorrhage, multi-organ failure (i.e., liver rupture, myocardial infarction, kidney failure), placental abruption, and heart disease later in life.^6–9^ Infants from preeclamptic pregnancies are at increased risk for intrauterine growth restriction (IUGR), fetal death, and premature delivery.^7^ While the definitive treatment for PE remains the timely delivery of the placenta and the baby, clinicians must carefully balance the urgent maternal need for delivery against the developmental benefits of prolonging the pregnancy for the fetus. Early identification of pregnancies at risk of PE enables tailored clinical management strategies^10^, including the initiation of low-dose aspirin before 16 weeks and enhanced monitoring, ultimately improving maternal and fetal outcomes.^11^ Recent advances in prenatal care, such as risk prediction models, diagnostic tests measuring angiogenic factors^12–15^, and the adoption of home monitoring technologies^16–18^, have significantly contributed to the management of PE. Despite these advancements, the lack of standardized and non-invasive testing continues to hinder the early identification of pregnancies at risk of PE. As such, there is an urgent need for consistent screening protocols to improve maternal and fetal outcomes.

### 2.2 Immune System Adaptations in Pregnancy and Preeclampsia

Pregnancy induces significant immune composition changes, promoting maternal tolerance and fetal development.^19^ In normotensive pregnancies, regulatory adaptations at the maternal-fetal interface involving immune cell function and immune cell numbers, promote maternal tolerance to the developing fetus, support fetal growth and maintain maternal health (Reviewed in:^19^). These changes include increased regulatory T cells (Tregs) recruited from maternal peripheral blood to the fetal-maternal interface, and a shift towards anti-inflammatory macrophages.^20^ However, in PE, these adaptions are disrupted, leading to a dominance of pro-inflammatory immune cells and a reduction in regulatory immune cells in peripheral blood.^20^ This imbalance in immune cell populations impedes maternal immune tolerance to the semi-allogenic fetus, and is thought to contribute to the pathogenesis of PE (Reviewed In:^2,21,22^). Monocytes, as precursors of tissue macrophages, are integral to the innate immune response, responsible for maintaining tissue integrity and responding to pathogens. In preeclamptic pregnancies, however, monocytes exhibit altered phenotype and functions, which contribute to the systemic inflammatory state characteristic of PE. Specifically, pro-inflammatory M1 macrophage activation increases, compounding the inflammatory response, leading to endothelial dysfunction and systemic inflammation.

Neutrophils, typically the first responders of the immune system, also display increased activation and adhesion in PE, leading to the release of reactive oxygen species and additional endothelial damage.^23^ Neutrophils serve as primary effectors in systemic inflammation by recruiting, activating, and reprogramming other immune cells, including dendritic cells, B cells, natural killer (NK) cells, CD4 and CD8 T cells, and mesenchymal stem cells. In PE, B cells also exhibit dysregulation, with altered antibody production that may further modify the placental immune environment.^22,24,25^

An imbalance in T cell subsets is another feature, characterized by an increased presence of pro-inflammatory T helper 1 (Th1) cells and a decrease in uterine and circulatory regulatory T cells (Tregs).^26^ Th1 cells release inflammatory cytokines, which further amplify the systemic inflammatory response in PE, while Tregs are essential for maintaining immune tolerance and suppressing excessive inflammation. This shift in the Th1/Treg ratio exacerbates the inflammatory conditions associated with PE.^21^ Moreover, a reduction in both the number and activity of NK is observed in PE.^27^ NK cells are crucial for immune surveillance and tolerance at the maternal-fetal interface, yet in PE, they display functional dysregulation within both maternal circulation and placental tissues. Collectively, these disruptions in the regulation of innate and adaptive immune cells and their cellular responses create a cytotoxic *in utero* environment, intensifying the pathological landscape of PE.^28^

### 2.3 Current Immune Markers and Limitations

In recent years, there has been a surge in research aimed at identifying predictive markers of PE, focusing on systemic inflammatory markers and immune cell signatures.^12,28–34^ Markers such as the neutrophil-to-lymphocyte ratio^29–31^ (NLR) and monocyte-to-lymphocyte ratio^35^ (MLR) are elevated in PE, associated with increased oxidative stress, endothelial dysfunction, and vascular damage. While promising, the utility of these immune cell ratios as a screening tool for PE are limited by sensitivity^34^, patient population applicability and comorbidities^32,33,36^, and predictive value for disease severity, restricting their effectiveness as screening tools.

### 2.4 Placental Interactions and Immune Cell Function

The complete extent of immune system dysregulation in PE is still not fully understood. The specific inflammatory pathways involved and the dynamic interactions between immune cells and other placental cells (e.g., trophoblasts and stromal cells) require further exploration. This complex interplay between immune cells and their interaction with placental cells likely forms a distinct cellular signature that may provide insights into the pathophysiology of PE and open avenues for potential diagnostic strategies. As trophoblasts play an essential role in early pregnancy by facilitating blastocyst attachment, invasion, migration, and endometrial remodelling, they are particularly susceptible to disruptions in the immunological microenvironment. Starting at 8 weeks of gestation, endovascular trophoblasts begin remodelling the spiral uterine arteries. The effectiveness of this placentation process is dependent on the ability of the extravillous trophoblasts to avoid detection by the maternal immune response.^37–39^ If maternal tolerance towards fetal components diminishes during this period, shallow placentation may occur, a condition frequently associated with PE.^40,41^

As pregnancy progresses, increased immune responses may exacerbate placental inflammation and lead to widespread systemic inflammation in the pregnant individual. Excessive activation of the maternal immune system can stimulate the release of proinflammatory cytokines and antiangiogenic factors within the fetoplacental unit and vascular endothelium.^42^ These molecules are critical mediators of intercellular communication but, in this context, amplify the immune response, further exacerbating systemic inflammation. The overactivation of maternal immune cells—including monocytes, macrophages, neutrophils, T cells, NK cells, and dendritic cells—against trophoblasts has been identified as an important mechanism triggering trophoblast apoptosis and cell death.^43^ This overactivity disrupts the remodelling of maternal uterine arteries, intensifying placental dysfunction. Abnormal proportions and functions of immune cells, including T cells, B cells, macrophages, and NK cells, create an immunological environment conducive to increased reactivity against trophoblasts.^21,22,39^ This heightened reactivity and immune dysfunction can be identified and measured by assessing immune cell counts and composition through advanced analytical tools.

### 2.5 DNA methylation as a Tool for Immune Composition Analysis

Advanced techniques such as flow cytometry, mass cytometry (CyTOF), single-cell RNA sequencing (scRNA-seq), and immunohistochemistry (IHC) have been used to accurately characterize immune cell activity and composition.^44,45^ Recently, DNA methylation (DNAm) cell deconvolution algorithms have emerged as a promising approach for identifying immune cell signatures, particularly in complex conditions like PE. Unlike traditional methods that require cell sorting, DNAm can infer immune cell proportions from bulk samples using cell-specific signatures and stable epigenetic modifications, making it especially useful in detecting subtle shifts in immune cell dynamics in PE.^46^

Most studies on DNAm changes during PE have focused on placental tissue^47^ or cord blood^48^, likely due to the historical emphasis on the placenta as the central driver of PE pathology and the accessibility of these tissues postpartum. However, maternal blood analysis is especially critical as it provides a direct and systemic view of immune-related changes, capturing cell-specific methylation patterns and proportions that could offer critical insights into maternal immune dynamics in PE. Reference-based algorithms^49^, which use DNAm data from sorted immune cell types obtained through flow cytometry, can estimate cell type proportions from bulk samples without requiring physical separation and sorting the immune cells. Given that both immune dysregulation and altered DNAm are both linked to PE^50^—and that DNAm plays a central role in immune cell development and function^51^—these shifts in cell proportions are inherently reflected in DNAm patterns associated with PE. Analyzing DNAm profiles of whole blood allows for the estimation of immune cell proportions, helping to pinpoint specific immune changes tied to PE. With a clinical need for reliable, pre-symptomatic PE prediction in both low- and high-risk pregnancies, profiling immune cell signatures through maternal blood sampling could be valuable for identifying and monitoring PE, supporting earlier intervention and improved patient management.

In this study, we used DNAm signatures from existing datasets to identify and characterize the immune cell composition in maternal blood samples from PE patients of different cohorts. This proof-of-concept study demonstrates that DNAm profiles in maternal blood can reflect specific immune cell shifts uniquely associated with PE, offering an approach to PE prediction and diagnosis. We hypothesized that the proportions of immune cell types would differ between preeclamptic and healthy pregnancies. Additionally, given the emerging research on DNAm changes throughout gestation, we further used DNAm profiled to investigate how immune cell composition might vary across different stages of pregnancy. By correlating DNAm signatures with altered immune cell proportions, this dual approach of identifying immune shifts across a normotensive gestational timeline and comparing normotensive and PE cases, provides deeper insights into the immunological and epigenetic landscape of PE, potentially leading to clinically viable method for early detection and monitoring of PE.

## 3.0 Methods

### 3.1 DNA methylation Sample Selection and Data Processing

This study analyzed DNAm datasets from maternal whole blood samples of pregnant individuals encompassing normotensive and preeclamptic pregnancies. A search query combining whole blood, pregnancy, and *Homo Sapiens* was employed, resulting in 34 studies (Figure 1). Out of the identified studies, five were further selected based on their profiling methods: Methylation profiling by array. All five studies involved samples described as either whole blood or peripheral whole blood. DNAm data were sourced from the Gene Expression Omnibus (GEO) database. To ensure high coverage, reproducibility, data quality and ease of analysis, the DNAm dataset screening was limited to the Illumina platform (i.e. Illumina 27K, 450K, and EPIC arrays). Two datasets, GSE37722 (HumanMethylation27 BeadChip arrays) and GSE192918 (Infinium MethylationEPIC BeadChip array), were selected for analysis. Both datasets were whole/peripheral blood samples. Only one study, GSE37722 included samples from normotensive (control, N=14) and preeclamptic pregnancies (N=14). To enhance the control sample size, data from GSE192918 were utilized, increasing the normotensive control group size to 24. In both datasets, the authors processed and normalized the raw data using GenomeStudio. Only the normalized data was available on GEO.

**Figure 1.**
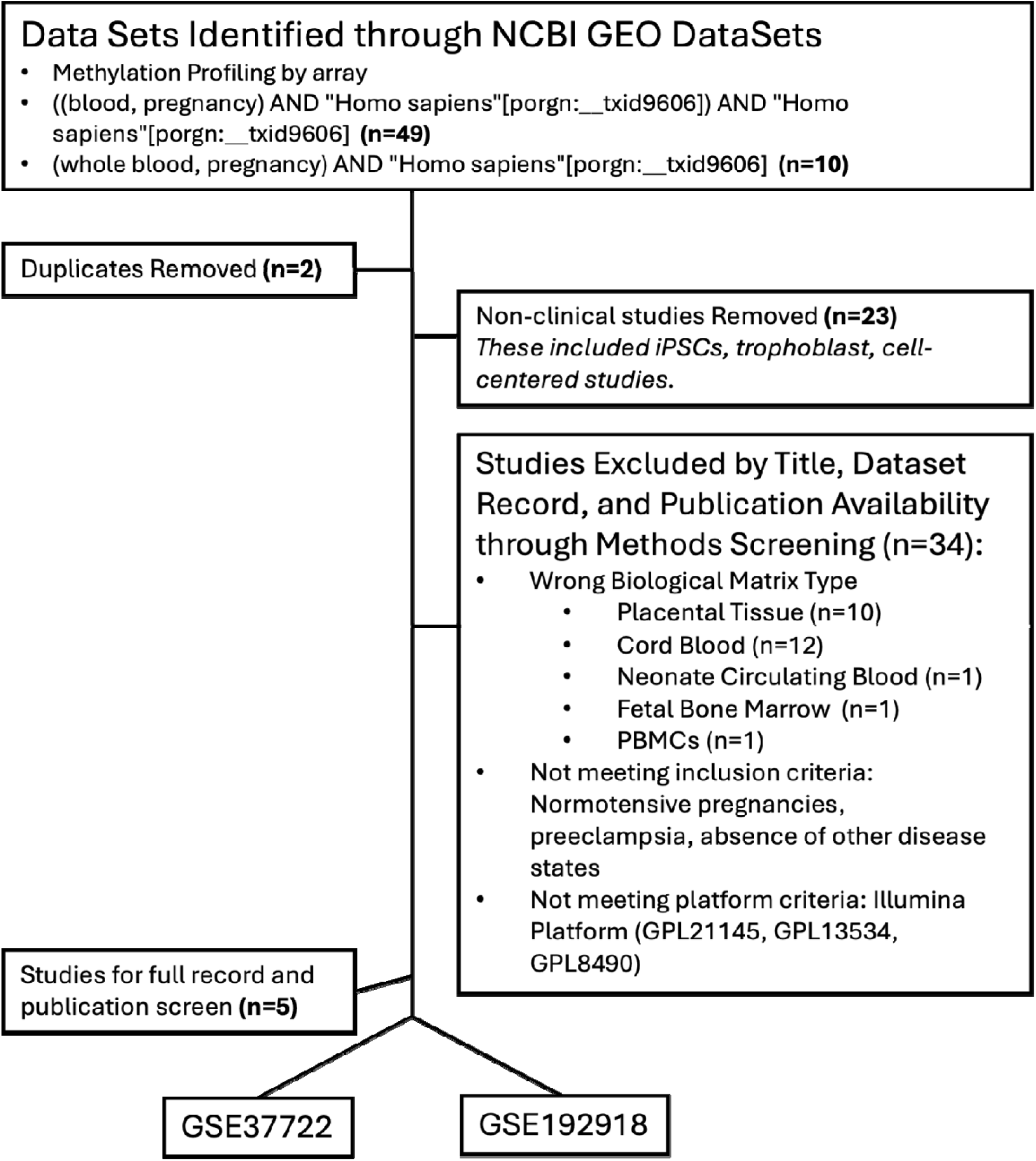
Flow chart of GEO dataset selection. The initial search yielded 59 studies. After removing two duplicates, 57 studies were screened for relevance based on the inclusion criteria, which included the use of human clinical samples (excluding human cells/stem cells/iPSCs). Further screening of the remaining 34 studies involved assessing biological matrix, normotensive pregnancies, preeclampsia, data sufficiency, absence of other disease states, and the use of the Illumina platform. Five studies met the criteria for deeper screening, of which two were selected for the final analysis.

DNAm data were systematically retrieved from GEO using the GEOquery package^52^ (version 2.66.0) within the R statistical environment (version 4.2.1). This procedure specifically targeted the Series Matrix files of the GSE37722 and GSE192918 datasets containing β-values, which are represent the ratio of the methylated probe intensity over the sum of methylated and unmethylated probe intensities. To compare normotensive and preeclamptic pregnancies, we identified common CpG sites between the datasets, as GSE37722 used the Illumina 27K array and GSE192918 used the EPIC array. This step ensured a consistent basis for analysis despite the different array platforms (24,128 probes remaining). Datasets were screened for missing data points (NAs), with any incomplete probes removed (20,404 probes remaining), prior to normalization. The datasets from GSE37722 and GSE192918 were merged, forming a master dataset. β-values were then normalized using quantile normalization (preprocessCore package, version 1.60.2).^53^ Although quantile normalization does not fully mitigate batch effects, it ensures comparability across datasets by aligning data distributions, enhancing the reliability of biological interpretations.^54^ Immune cell deconvolution was then performed using the Epigenetic Dissection of Intra-Sample Heterogeneity (EpiDISH) algorithm^55,56^ to infer cellular composition from DNAm profiles.

### 3.2 Cell-deconvolution for Leukocyte Composition Estimation

EpiDISH is a computational algorithm used for the deconvolution of epigenetic data to estimate the proportion of different cell types (epithelial cells, fibroblast cells, and seven immune cell subtypes: neutrophils, eosinophils, monocytes, CD4+ and CD8+ T cells, B cells, and Natural Killer cells) in a mixed cell population found in whole blood, generic epithelial tissue, and breast tissue^56–60^. The *EpiDISH* package^55,56^ (version 2.14.1) utilizes a reference-based approach to deconvolute DNAm data into estimates of cellular proportions, leveraging known reference DNAm profiles of cell types of interest. The merged dataset was analyzed using the robust partial correlations (RPC) method. The reference matrix *centDHSbloodDMC.m*^55,61^ for blood cell types was applied to estimate immune cell-type fractions in the samples of interest. The immune cell proportion data was compared between ‘Normotensive’ (which included GSM identifiers from both the GSE37722 and GSE192918 datasets) and ‘Preeclamptic’ pregnancies (which included only GSM identifiers from the GSE37722 dataset). The data analyzed in this study were obtained from publicly available datasets in the Gene Expression Omnibus (GEO), specifically under accession numbers GSE37722 and GSE192918. The processed data and analysis scripts used in this study are openly available in the GitHub repository at https://github.com/WilsonPregnancyLab/ImmuneCellComposition.

#### Changes in immune cell composition across gestation

The datasets GSE192918 and GSE37722 were both used to independently investigate immune cell composition across different stages of pregnancy. For each dataset, the series matrix was individually extracted, and sample IDs were identified and matched to specific gestational timepoints. These stages were categorized as ‘early-pregnancy’, ‘mid-pregnancy’, ‘at delivery’ and ‘post-partum’. GSE192918 defined these stages as follows: early-pregnancy (10-14 weeks), mid-pregnancy (24-28 weeks), at delivery (38-40 weeks), and post-partum (10 month post-delivery). While GSE37722 also included data for normotensive pregnancy stages (i.e., early, mid, delivery, postpartum), the exact timings of these stages were not explicitly defined in the GEO database. Subsequent processing involved quantile normalization of both datasets, application of the EpiDISH algorithm to estimate immune cell fractions, and stratification of these estimates by gestational timepoint.

### 3.3 Data Visualization and Statistics

#### Comparing Normotensive and Preeclamptic Pregnancies

All statistical analyses were conducted within the R statistical environment (version 4.2.1). As the datasets did not meet the assumptions of normality and equal variance, the Wilcoxon rank-sum test was used to determine significant difference in cell-type proportions and DNAm levels between normotensive and preeclamptic pregnancy groups. The Kolmogorov-Smirnoff test was used to determine differences in the overall distribution of cell type composition between normotensive and preeclamptic pregnancies. A p-value of ≤ 0.05 was considered statistically significant.

#### Changes in immune cell composition across gestation

Linear modelling was used to assess changes in the proportion of immune cell subtypes at different time points across gestation in both GSE192918 and GSE37722. For each immune cell subtype, a linear model was constructed using the limma package^62^ (version 3.54.2) in R. Contrasts were applied to compare the estimated fraction of each immune cell subtype between each pairwise combination of time points. A false discovery rate (FDR)-corrected p-value was calculated for each comparison. An FDR cut-off of 0.05 was used as a threshold for significance. Plots for this data were produced with the ggplot2 library (version 3.4.3) in R. Additionally, an Anderson-Darling all pairs comparison test was used to assess differences in the overall distribution of cell type proportions across timepoints. All pairwise combinations of timepoints were tested. A p-value of ≤ 0.05 was considered statistically significant.

## 4.0 Results

### Differential proportion of monocytes in preeclamptic versus normotensive pregnancies

Immune cell proportion of neutrophils, eosinophils, NK cells, B cells, and both T cell subtypes, did not exhibit statistically significant changes between normotensive and preeclamptic pregnancy groups (Figure 2). The proportion of monocytes was significantly reduced in the PE group.

**Figure 2.**
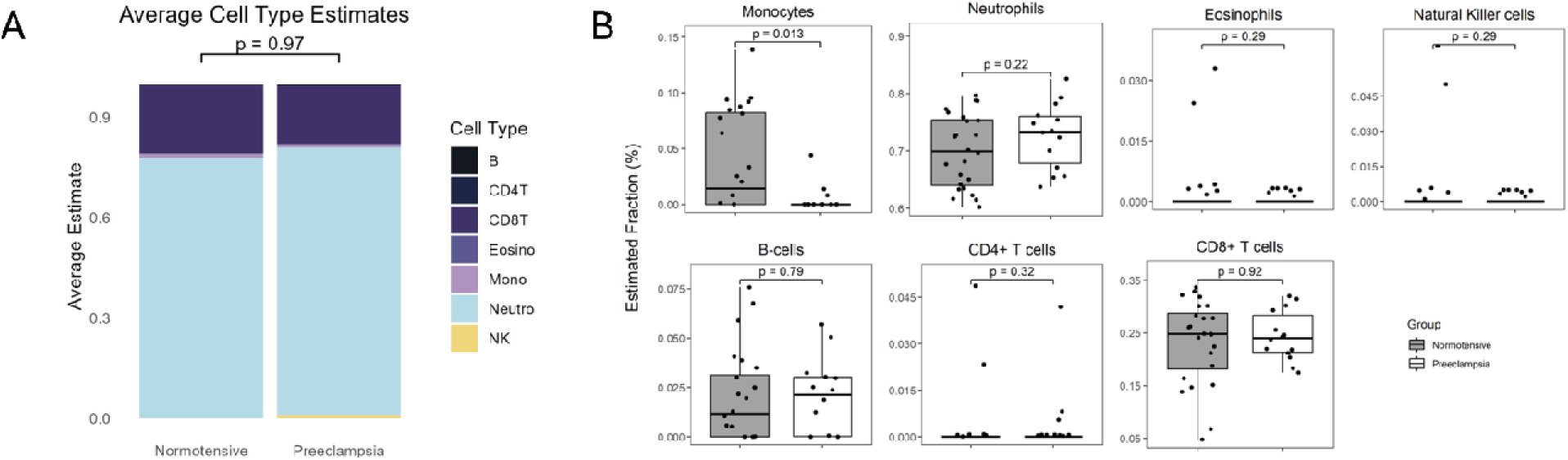
(A) Average Estimated Cell Type Distribution and (B) Immune cell proportion between normotensive (grey, control) and preeclamptic (white) pregnancies. EpiDISH was used to deconvolute GSE37722 (N=14 control, N=14 preeclamptic pregnancies) and GSE192918 (N=10 control) to identify changes in the proportion of monocytes, neutrophils, eosinophils, natural killer cells, B-cells, CD4+ T cells and CD8+ T cells. All data is presented as mean. A Kolmogorov-Smirnoff test was used to assess differences in the overall distributions, while a Wilcoxon test was conducted for each cell type. A p-value of ≤ 0.05 was considered statistically significant.

### Immune cell composition varies with gestational age, with the most pronounced changes occurring in the post-partum period

In our analysis of the gestational age across datasets, significant changes were observed in the immune cell profiles between the delivery and post-partum periods (Figures 3, 4, 5). Although there were no significant differences between trimesters in either dataset, the greatest changes in immune cell profiles were observed postpartum. In dataset GSE37722, which included blood samples taken within 24 hours of delivery, significant changes in immune cell profiles were observed between the delivery and post-partum periods (Figure 3).

**Figure 3.**
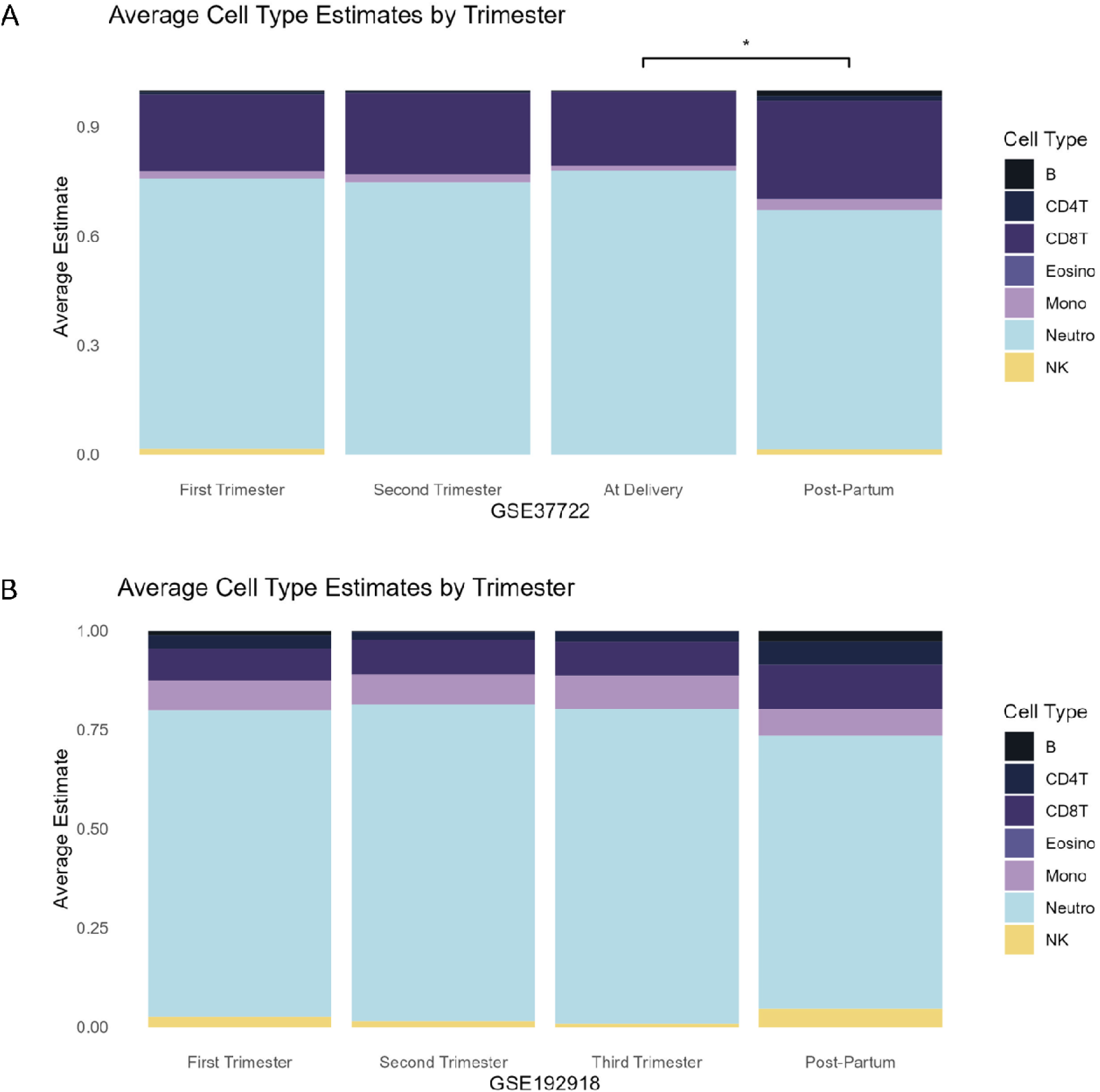
Average Estimated Cell Type Distribution across gestation for (A) GSE37722 and (B) GSE192918. EpiDISH was used to deconvolute and identify the proportions of monocytes, neutrophils, eosinophils, natural killer cells, B-cells, CD4+ T cells and CD8+ T cells across ‘early-pregnancy’, ‘mid-pregnancy’, ‘at delivery’ and ‘post-partum’. While GSE37722 included data for normotensive pregnancy stages, the exact timings of these stages were not explicitly defined in the GEO database. Differences in distribution across gestational were assessed using an Anderson-Darling all pairs comparison test. A p-value of ≤ 0.05 was considered statistically significant.

Specifically, there was a notable increase of monocytes, B-cells, CD4+ T cells, and CD8+ T cells (Figure 4). B-cells were significantly increased when each pregnancy stage (early, mid, and at delivery) was individually compared to the post-partum period. Interestingly, in a similar comparison, the neutrophil proportion was significantly reduced in the postpartum period. In contrast, natural killer (NK) cells showed a higher proportion during early pregnancy compared to middle or at delivery. Notably, NK cell proportions returned to early pregnancy levels in the post-partum period.

**Figure 4.**
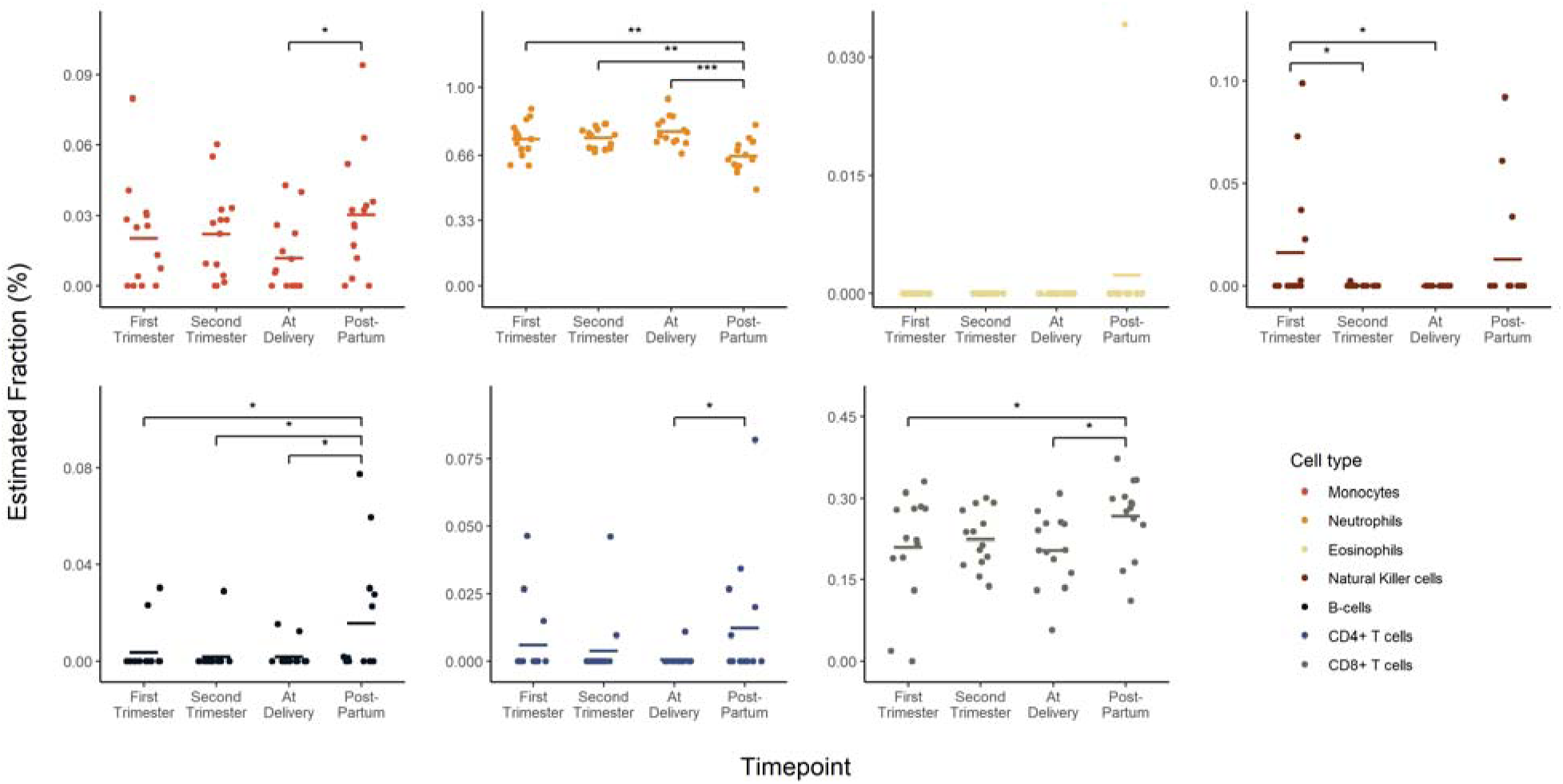
Immune cell composition across gestational age within each GSE dataset (GSE37722). EpiDISH was used to deconvolute and identify the proportions of monocytes, neutrophils, eosinophils, natural killer cells, B-cells, CD4+ T cells and CD8+ T cells across ‘early-pregnancy’, ‘mid-pregnancy’, ‘at delivery’ and ‘post-partum’. While GSE37722 included data for normotensive pregnancy stages, the exact timings of these stages were not explicitly defined in the GEO database. All data is presented as mean. A linear model was conducted for each cell type. An FDR-adjusted p-value of ≤ 0.05 was considered statistically significant.

In our analysis of the GSE192918 dataset, similar to GSE37722, significant changes were noted in the immune cell profiles from delivery to the post-partum period. Notably, there was a significant induction of NK cells, B-cells, CD4+ T cells, and CD8+ T cells (Figure 5). Specifically, B-cells and CD8+ T cells showed significant increases when comparing each pregnancy stage (early, mid, and at delivery) to the post-partum period, whereas neutrophils significantly decreased. There were no changes in monocytes across gestation or in the post-partum period.

**Figure 5.**
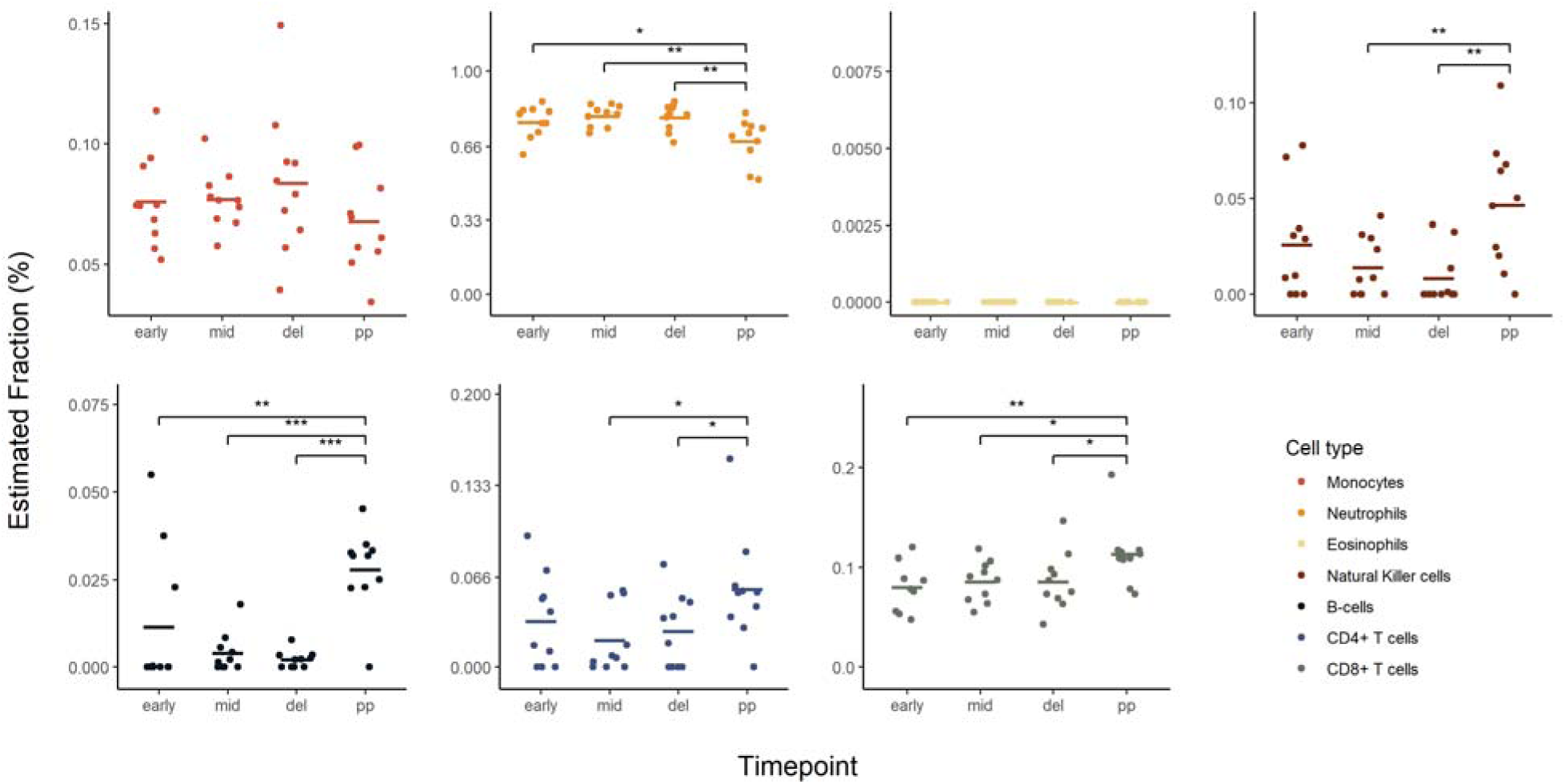
Immune cell composition across gestational age within each GSE dataset (GSE192918). EpiDISH was used to deconvolute and identify the proportions of Neutrophils, Monocytes, Natural Killer cells, Eosinophils, B-cells, CD4+ T cells and CD8+ T cells across ‘early-pregnancy’, ‘mid-pregnancy’, ‘at delivery’ and ‘post-partum’. GSE192918 defined these stages as follows: early-pregnancy (10-14 weeks), mid-pregnancy (24-28 weeks), at delivery (38-40 weeks), and post-partum (10 month post-delivery). While GSE37722 also included data for normotensive pregnancy stages, the exact timings of these stages were not explicitly defined in the GEO database. All data are presented is mean ± SEM. A linear model was conducted for each cell type. An FDR-adjusted p-value of ≤ 0.05 was considered statistically significant.

## 5.0 Discussion

Numerous studies have explored the relationship between leukocyte counts and PE to better understand the underlying immune dysregulation (Reviewed in: ^29^). Our findings provide novel insights by leveraging DNAm signatures to characterize immune cell proportions in maternal blood, offering an efficient alternative to more labor-intensive techniques like flow cytometry. While overall immune cell composition—such as neutrophils, eosinophils, NK cells, and T cell subtypes—did not differ significantly between normotensive and preeclamptic pregnancies, we observed a notable reduction in monocyte proportions in the PE group. This finding is consistent with previous studies suggesting that altered monocyte behaviour (i.e., reduced trafficking or functional impairment), rather than sheer number, plays a key role in the inflammatory profile and pathogenesis of PE. Differentiating between increased leukocyte counts and specific monocyte behaviour in PE us critical, as evidence increasingly shows that monocyte counts in PE either remain stable or show a downward trend, in contrast to general leukocyte increases commonly seen in hypertensive pregnancies.^63–66^

Monocytes, comprising approximately 3-8% of peripheral blood leucocytes, are essential components of the innate immune system. They serve as progenitor cells with the ability to differentiate into macrophages and dendritic cells, contributing to both immune surveillance and tissue remodelling^67,68^. In a typical pregnancy, there is an expected increase in innate immune cell activation in circulation, reflected by elevated monocytes and granulocytes counts, peaking between 29-36 weeks, and increased production of pro-inflammatory cytokines (e.g., IL1b, IL6, IL8).^69,70^ PE, however, represents an exaggerated state of immune activation where monocytes exhibit heightened activation and dysfunctional surface antigen expression, impaired vascular adherence, and reduced immunosuppressive capacities, which likely contribute to the systemic inflammation in PE.^43^ Sacks et al. (1998)^63^ demonstrated a decrease in CD14 expression on monocytes and granulocytes in PE, suggesting a functional shift in monocytes rather than a change in overall count. Melgert et al. (2012)^64^ and Al-ofi et al. (2012)^65^ corroborated that, despite a general increase in leukocyte counts, monocyte numbers themselves did not differ between normotensive and preeclamptic pregnancies. Notably, Sacks et al. based PE diagnoses on the American College of Obstetricians and Gynecologists (ACOG) criteria^71^, while Melgert and Al-ofi used International Society for the Study of Hypertension in Pregnancy (ISSHP) definitions^72^—these differing criteria, along with the inherent heterogeneity of PE^73^, could partly explain the variability across studies. These definitional differences highlight the complexity of studying PE, as different diagnostic criteria may lead to variability in the severity and characteristics of PE captured in each study. Despite these differences, the studies consistently indicate that it is monocyte functionality, rather than count, that plays a significant role in PE pathophysiology. In line with this, Wang et al (2019), concluded that compared to absolute cell counts, monocytes-to-lymphocytes ratios offer a more effective indication of clinical assessments, disease severity evaluation, and prognosis evaluation of PE.^35^ However, other studies exploring the MLR have reported mixed findings. While some studies by Bektaş et al. (2019)^74^ support Wang et al. (2019)^35^ and found MLR to be elevated in PE, Seyhanli et al. (2024)^75^ reported no significant effect, and Cui et al. (2023)^36^ suggested that all inflammation indices were lower in PE patients. This inconsistency in literature highlights the need for alternative approaches that can offer a more consistent and precise understanding of immune alterations in PE.

Whereas studies investigating absolute immune cell counts^63–65^ use flow cytometry to directly evaluate cell surface markers and monocyte activation, our approach applies a DNAm-based reference algorithm (EpiDISH) to infer immune cell proportions and revealing a decrease in monocyte proportions in PE. Although prior studies found either stable or slightly declining monocyte counts in PE, our DNAm-based approach detected a statistically significant reduction. This difference could reflect the sensitivity of DNAm signatures and deconvolution in detecting subtle proportional changes that flow cytometry may not capture when analyzing absolute counts. Flow cytometry is a well-established cell-based technique that provides high-resolution data on cell surface markers and immune cell subsets but has limitations, including the need for fresh samples, labor-intensive processing, and restricted analysis of only a limited number of markers simultaneously. In contrast, DNAm-based deconvolution, offers a computational approach, allowing immune cell estimation by detecting unique, cell-specific epigenetic signatures. These signatures provide insights into immune composition that are not solely dependent on surface antigen expression, thus allowing for broader cell-type identification in bulk, complex, or archived tissue samples.^76^ Despite these advantages, DNAm deconvolution has limitations. It relies on reference matrices to estimate cell types, inherently providing estimates rather than direct measurements, potentially introducing bias and inaccuracies, especially when the reference matrices may not perfectly reflect the biological characteristics of the study population.

The reference matrix used in this analysis (EpiDISH: centDHSbloodDMC.m) was derived from six healthy male blood donors (aged 38±13.6 years)^61^, correlating flow cytometry-immunophenotyped PBMCs with DNAm data from the Illumina 450k platform (GSE35069). While this matrix has been validated across multiple biological matrices for immune cell estimations, it may not fully capture the unique immunological and hormonal dynamics associated with biological sex^77–79^ or pregnancy, particularly under pathological conditions such as PE. However, it is important to emphasize that, despite these limitations, our results remain robust. Both the normotensive and PE groups were analyzed using the same male-derived reference matrix, so any biases introduced by this reference would affect both groups equally, preserving the validity of relative comparisons. This approach ensures that, while absolute monocyte proportions may be influenced by the male-derived matrix, the observed reduction in monocyte proportions in the PE group compared to the normotensive group remains significant and reliable. Future work could address the assumption that immune cell proportions estimated from a male reference matrix generalize well to female samples, as current literature lacks a direct assessment of this assumption.

A key limitation of DNAm-based reference algorithms, including EpiDISH, is their dependence on such reference matrices, which are typically developed by correlating DNAm data with flow cytometry. While flow cytometry can differentiate specific monocyte subsets, such as classical (CD14+CD16−) and non-classical (CD14lowCD16+) monocytes, as well as monocyte-derivatives like dendritic cells and macrophages, few reference matrices currently include these subsets. This limitation restricts DNAm deconvolution’s ability to capture the full complexity of immune cell profiles in PE, where subset-specific responses are crucial to understanding disease progression.^80^ In PE, inflammatory monocyte subsets like non-classical monocytes contribute to vascular dysfunction, potentially increasing inflammation at the maternal-fetal interface.^81^ Macrophages in PE also show an imbalance between M1 (pro-inflammatory) and M2 (anti-inflammatory) types, with increased M1 macrophages impairing trophoblast migration and vascular remodeling (Reviewed in: ^70,82^). Dendritic cells, which are typically inactive near the placenta during healthy pregnancies, become improperly activated in PE, compounding the inflammatory environment. DNAm patterns among these monocyte subsets^81,83^ and their derivatives^61,83,84^ are known but not yet integrated into current EpiDISH matrices, leaving it unclear if the observed monocyte decrease is due to differentiation into macrophages or other changes in subset activation. Together, these data and this complexity suggests that monocyte counts alone may not distinguish between healthy and PE pregnancies. Instead, a combination of monocyte count, activation state, and phenotypic changes likely underpins the immune dysregulation seen in PE. Future research should refine reference matrices to better capture the immune dynamics of pregnancy and develop DNAm tools for differentiating monocyte subsets in epigenetic analyses.

Monocyte counts are also influenced by gestational age and critical windows within pregnancy, contributing shifts in total cell counts, subset proportions, and differentiation patterns (e.g., macrophages, dendritic cells, T cells) across stages of pregnancy, particularly around parturition. The GSE37722 Illumina 27k methylation array data used in this study was extracted from blood samples collected within 24h of delivery, a critical period marked by distinct immune responses.^85^ During parturition, immune activation includes a shift in monocyte activation not seen earlier in pregnancy.^86,87^ Studies report lower non-classical monocytes (CD14lowCD16+) and higher intermediate monocytes (CD14+CD16+)^70,88^ in laboring patients, highlighting the importance of gestational timing when analyzing immune responses in pregnancy. Interestingly, immune composition alterations are observed throughout the third trimester and intro the postpartum period,^64,70^ reflecting the dynamic nature of immune modulation from early pregnancy through delivery.

Gestational adaptation of the maternal immune system begins soon after fertilization and continues throughout pregnancy, with a return to a pre-pregnancy state after delivery and lactation.^19^ To examine immune composition changes across gestational timepoints, we analyzed datasets GSE37722 and GSE192918. While the role of DNAm in regulating maternal physiology during pregnancy and postpartum is still not fully understood, our study provides new insights into these dynamics. At delivery, genome-wide DNAm profiles in maternal leukocyte DNA were more methylated in PE cases than in normotensive controls, particularly at promotor CpG sites, suggesting that many genes may be repressed or downregulated, potentially contributing to immune dysregulation.^85^ Analysis of dataset GSE192918^89^ revealed that 61.63% of CpG sites displayed subtle DNAm changes during pregnancy, with more significant changes occurring postpartum. Notably, the second and third highest concentrations of altered DNAm were observed between the first and second trimesters, likely reflecting key phases of embryonic development and organogenesis. Pathway analyses from GSE192918 highlighted several immune-related pathways, including IL-15 production, Fcγ receptor-mediated phagocytosis in macrophages and monocytes, TREM1 signaling, and IL-7 signaling, suggesting that DNAm changes may shape the maternal immune landscape.^89^ This potential reshaping of the immune environment is further supported by our findings, which showed significant shifts in immune cell composition—particularly in neutrophils, CD8+ T cells, and B cells—between pregnancy and the postpartum period.

Immune cell analysis in both datasets revealed that the most significant difference in cell counts occurred between pregnancy (early, mid, at delivery) and the postpartum period. Neutrophils, CD8+ T cells, and B cells showed substantial shifts, especially during the transition from pregnancy to postpartum. Neutrophil counts, which varied by gestational age, were consistently lower postpartum. These shifts likely represent the immune system’s recovery from the pregnancy-associated immunotolerant state and a reactivation of systemic responses. Although we initially hypothesize trimester-specific changes, immune cell composition remained relatively stable across trimesters. This finding contrasts with earlier reports of trimester-specific immune shifts in pregnancy^19,90,91^, emphasizing the complexity of immune modulation at the maternal-fetal interface. However, the postpartum period marked significant changes, with increased CD8+ T cells, B cells, and monocytes, which may indicate a rebound of systemic immunity following the immunosuppressive state of pregnancy. While symptoms of PE usually end upon delivery of the placenta, inflammation may persist into the mother’s postpartum period (Reviewed In: ^92^).

Discrepancies between GSE37722 and GSE192918, particularly regarding NK cell and monocyte shifts, could be attributed to small sample size and differences in population characteristics. For example, GSE37722 included only first-time mothers of European descent^85^, while GSE192918 included 10 women with uncomplicated pregnancies^89^. Although these datasets controlled for key variables such as smoking status, BMI, and age^85,89^, other factors such as ethnicity and environmental exposures may still influence DNAm. Additionally, differences in platforms used to measure DNAm introduce challenges for cross-study comparisons. The limited clinical data available for these datasets further complicates interpretations, making it difficult to fully understand the factors influencing variations in immune profiles. Both datasets, however, emphasize the dynamic nature of immune adaptation throughout pregnancy and into the postpartum period.

The heterogeneity of PE itself likely contributes to these variations, as PE can be divided into early-onset PE (early-PE), diagnosed at or before 34 weeks of gestation, and late-onset PE (late-PE), diagnosed after 34 weeks.^9,93^ These subtypes involve distinct pathophysiological processes. Early-PE is commonly linked to placental dysfunction and often presents with intrauterine growth restriction (IUGR), while late-PE has no consensus in etiology.^94,95^ It is hypothesized to be driven by maternal factors or later-developing placental issues. Immune dysregulation plays a role in both subtypes but likely manifests differently in each.^96^ Transcriptional and epigenetic analyses are revealing new molecular phenotypes of PE that may not align directly with traditional clinical classification. Leavey et al. (2018, 2019) identified two additional transcriptional subtypes of PE - *canonical* and *immunological* - distinguished by unique gene expression profiles and epigenetic patterns in placental tissue.^97,98^ These molecular phenotypes along with observed variations in placental immune cell composition suggest that there are additional complexities and mechanisms that extend beyond clinical definitions of PE.

Although DNAm changes may have the potential to differentiate between early-PE, late-PE, and healthy pregnancies^97,99–101^, our study is limited by the lack of phenotypic data within the datasets used^85,89^. This absence of specific subtype information means that we cannot make determinations about whether the observed DNAm changes are specific to a particular subset of PE or reflect general PE-related immune dysregulation. Furthermore, publicly available datasets like GSE192918 and GSE37722 are constrained by small sample sizes, reducing statistical power to detect subtle changes, especially in heterogeneous conditions like PE.^102^ Data availability, publication delays, and technical variability across studies further hinder progress in rapidly evolving fields like epigenetics and immunology. Future studies with comprehensive phenotypic data, including detailed information on PE onset and subtype, would enable a more accurate assessment of DNAm’s ability to distinguish between early-PE, late-PE, and healthy pregnancies, thereby providing a clearer understanding of these epigenetic signatures.^102^

Our findings also indicate that DNAm can serve as a tool to infer immune cell composition and detect immune system activation across pregnancy stages. While most previous studies have focused on DNAm changes in fetal-derived tissues or cell-free placental DNA^47,48,97,98,101,102^ in maternal plasma, our study demonstrates that maternal leukocyte DNA provides valuable insights into immune changes during pregnancy and PE.^85^ The observed DNAm changes in PE, especially increased DNAm in maternal leukocytes, suggest that epigenetic modifications may play a role in the pathogenesis of PE, potentially by regulating key immune-related genes.

Future research should aim for larger, more diverse cohorts and harmonized methodologies to enhance our understanding of the relationship between DNAm, immune cell dynamics, and PE development. A critical component in advancing this field is the development of more robust reference matrices. Specifically, reference matrices derived from female samples, with finer resolution of immune cell subsets through advanced flow cytometry, would better capture the unique immune landscape of pregnancy and PE. Additionally, instead of matrices designed solely to address specific scientific questions, comprehensive, large-scale reference matrices incorporating multiple factors and diverse sampling sites would enable more accurate and personalized assessments of immune profiles. Researchers are encouraged to adhere to the four-step checklist for biological data deposition by Wilson et al. (2021)^103^, ensuring that data shared across specialist and general repositories adheres to the FAIR guiding principles—Findable, Accessible, Interpretable, and Reusable. This approach will promote consistency, transparency, and broader utility in future studies.

## 6.0 Conclusion

This study demonstrates the potential of DNAm profiling as an insightful tool for understanding immune cell dynamics throughout pregnancy and PE. Our findings suggest significant differences in DNAm patterns between preeclamptic and normotensive pregnancies, which may reflect altered immune responses contributing to the disease’s pathogenesis. Notably, we observed significant differences in monocyte proportions between PE and normotensive pregnancies, adding to the growing body of evidence that immune system dysregulation plays a central role in PE pathogenesis. Significant immune cell shifts were observed during the transition from pregnancy to the postpartum period, reinforcing the importance of longitudinal sampling to fully capture immune changes associated with pregnancy complications. Recent evidence suggests that a multitude of factors—including genetic predisposition, environmental exposures, vascular dysfunction, placental abnormalities, and immune system activation—contribute to the onset and distinct clinical presentations of PE. Our study supports this multifactorial view, with epigenetic changes likely reflecting and affecting these diverse influences. The ability to use DNAm profiles to track immune cell changes across pregnancy and into the postpartum period offers promise for developing more refined diagnostic and monitoring tools for complications like PE. By incorporating longitudinal data and postpartum analysis, future research can better capture the full spectrum of immune dynamics and their potential long-term effects on maternal and fetal health. Despite these advances, limitations related to small sample sizes, technical variability, and inconsistent clinical data must be addressed. Future research with larger, more diverse cohorts and harmonized methodologies will be essential to deepen our understanding of the complex interplay between epigenetic regulation and immune function in PE. Adopting FAIR data principles will also ensure that future findings are transparent, reproducible, and broadly accessible, helping to advance maternal health research on a global scale.

## 7.0 Data Accessibility Statement

The data analyzed in this study were obtained from publicly available datasets in the Gene Expression Omnibus (GEO), specifically under accession numbers GSE37722 and GSE192918. The processed data and analysis scripts used in this study are openly available in the GitHub repository at https://github.com/WilsonPregnancyLab/ImmuneCellComposition.

The repository includes the R scripts for immune cell proportion estimation using the *EpiDISH* package, statistical analyses, and visualizations. Researchers are encouraged to access, reproduce, and build upon our work using the resources provided. For additional inquiries or clarifications, please contact the corresponding author.

## Notes

https://www.ncbi.nlm.nih.gov/geo/query/acc.cgi

### Competing Interest Statement

The authors have declared no competing interest.

